# Estimating the fundamental niche: accounting for the uneven availability of existing climates in the calibration area

**DOI:** 10.1101/2021.01.25.428165

**Authors:** L. Jiménez, J. Soberón

**Affiliations:** School of Life Sciences, University of Hawai’i at Mānoa, 2538 McCarthy Mall, Honolulu, HI 96822, USA; Biodiversity Institute, University of Kansas, 1345 Jayhawk Blvd, Lawrence, KS 66045, USA

**Keywords:** realized niche, environmental space, presence data, weighted distribution, dispersal

## Abstract

Studies that question important conceptual and methodological aspects of the field of ecological niche modeling (and species distribution modeling) have cast doubts on whether it is possible to estimate the fundamental niche of a species using presence-only data. The main limitation in niche estimation is that presence data come from the realized niche, which is only a subset of the fundamental niche. Most of the existing methods lack the ability to overcome this limitation and therefore fit niches that resemble the realized niche. To obtain a more accurate estimate of the fundamental niche, we propose using the geographic region that is accessible to a species (based on its dispersal ability) to determine a sampling distribution, in environmental space, from which we can quantify the likelihood of observing a particular environmental combination in a sample of presence points. We incorporate this sampling distribution into a multivariate normal model (i.e., a Mahalanobis distance model) by creating a weight function that modifies the probabilities of observing different environmental combinations in a sample of presences. This modification accounts for the uneven availability of environmental conditions. We show that the parameters of this weighted-normal model can be approximated with a maximum likelihood estimation approach; and then used to draw ellipsoids (confidence regions) that represent the fundamental niche of the species. We illustrate the application of our model with two worked examples. First, we use presence data for an invasive species and an accessible area that includes only its native range to evaluate whether the fitted model predicts confirmed establishments of the species outside its native range. Second, we use presence data for closely related species with known accessible areas to demonstrate how the different dispersal abilities of the species constrain a classic Mahalanobis distance model. Overall, we show that accounting for the distribution of environmental conditions that are accessible to a species indeed affects the estimation of the ellipsoids used to model its fundamental niche.

## 1 Introduction

In recent years, there has been substantial progress in the fields of ecological niche modeling (ENM) and species distribution modeling (SDM) (Guisan et al., 2013). However, there is still debate about what aspects of the niche are estimated by these methods (Jiménez-valverde et al., 2008; Lobo, 2008; Warren, 2012). Specifically, conventional ENM/SDM approaches that are based on presence-only data estimate objects that, in the case of ENM, are between the realized and the fundamental niches (Peterson et al., 2011), or, in the case of SDM, are between the actual and potential distributions of the species (Jiménez-valverde et al., 2008). Here, we propose a model to estimating fundamental niches that fits biologically realistic shapes and quantifies the availability of climates that are accessible for the species, as a way to account for biased presence samples.

The distinction between the fundamental and the realized niche, as proposed by Hutchinson (1957), is essential to understand what kind of objects are being estimated by the different correlative statistical models used in ENM/SDM. The fundamental niche of a species is the set of environmental conditions where, in the absence of biotic interactions, the population growth rate is positive (Peterson et al., 2011). The realized niche is a subset of the fundamental niche that is determined by abiotic factors (environmental conditions), biotic factors, and dispersal limitations (Soberón, 2007). Estimating the fundamental niche of a species is of particular importance when using the estimated niche to model species distributions at other times or in different regions, such as when using ENM/SDM to predict the effects of climate change or the spread of invasive species (Tingley et al., 2014). However, determining the complete fundamental niche of a species is also substantially more difficult than estimating the realized niche. It has being suggested that, in principle, we should not expect models to predict the fundamental niche, unless some supplementary information on it (allowing extrapolation beyond the realized niche) is used to fit the models (Qiao et al., 2018). In that sense, some approaches have incorporated experimental data on the physiology of the species to their models as a source of information from unoccupied regions of the fundamental niche (Hoogenboom & Connolly, 2009; Jiménez et al., 2019). Additionally, Booth (2017) has suggested that, for well-known tree species, different types of occurrences beyond their natural distributions (e.g., commercial forestry trials where competition is reduced) can be used as supplementary information on the species climatic requirements.

The relationship between modeling niches and modeling geographic distributions is mediated by Hutchinson’s duality (Colwell & Rangel, 2009), which is the relationship between environmental and geographic spaces. With the right resolution, a discrete set of geographic coordinates can be made to have a one-to-one relationship with a discrete set of environmental vectors (Aspinall & Lees, 1994; Soberon & Nakamura, 2009). This fundamental correspondence allows us to move back and forth between modeling niches and modeling geographic distributions. As a consequence of Hutchinson’s duality (Colwell & Rangel, 2009), and because species presence data come only from areas currently occupied by a species, a sample of presence records may not reflect all the environmental potentiality of a species (Jiménez-valverde et al., 2008; Lobo, 2008). Therefore, we should acknowledge that a correlative model based solely on presence data will be constrained by the imposed limitations of the set of environments where the species can be observed (Owens et al., 2020), will probably approximate the realized niche (Soberon & Nakamura, 2009). Failing to acknowledge and (somehow) include this constraint in a model leads to severe uncertainties and drawbacks when using this models to make predictions (Lobo, 2016).

Species presence/absence data are often spatially biased and noisy. We work under the assumption that a cleaning and preparation process to resolve errors precedes the application of ENM/SDM methods. Several techniques have been developed for resolving some of the common types of problems in presence data (Chapman, 2005; Cobos et al., 2018), such as a lack of accuracy in the reported coordinates, nomenclatural and taxonomic errors, and the presence of geographic or environmental outliers. However, other types of bias remain in presence data. We focus on the implicit bias created when defining the relevant spatial region for a study. Selecting different study regions produces different sampling universes, and the definition of the sampling region is an important part of the specification of a model. Here, we explicitly define an “M” hypothesis to set the sampling universe as all the sites where we can observe the species as present (Barve et al., 2011).

Under the “biotic-abiotic-movement” (BAM) framework (Peterson et al., 2011), the region M contains all the sites that a species is hypothesized to have been able to reach from some past time (via dispersal and migration). In geography (G-space), M is usually a connected and continuous set (i.e., a single polygon), so we make use of the Hutchinson’s duality to perform practical computations. This set is first converted into a discrete grid of coordinates, and then the values for environmental variables at these coordinates are used to build its representation in environmental space (E-space). Because the fundamental niche is defined in E-space, it makes sense to represent our data (M hypothesis and presences) and models in E-space.

There are two things that we need to consider regarding the relationship among M and the fundamental niche. First, not all sites in M have the environmental conditions needed to sustain viable populations, and it is known that some sites inside M could be sink populations (Barve et al., 2011). Second, Hutchinson’s duality permits the establishment of a one-to-one relationship between coordinates in G-space and multidimensional vectors in E-space, but this relationship does not preserve distance: nearby elements in G-space are not necessarily nearby in E-space, and vice versa. This creates a serious, and mostly ignored problem: uniform, random sampling in G-space does not imply uniform, random sampling in E-space. If the goal of a niche model is to get estimate the fundamental niche using presence-only data, the empirical distribution of M in E-space should therefore be incorporated in the statistical model to account for the uneven distribution of available sampling points.

In a previous contribution (Jiménez et al., 2019), we proposed a Bayesian argument for combining correlative techniques with information from physiological experiments to approximate to the fundamental niche of a species. However, in that work, we disregarded the problem of heterogeneity of E-space. That is why here we consider that event of observing an environmental combination ***x*** ∈ ℝ^*d*^ (where the species was recorded as present) has a non-uniform probability of being included in the sample. In a typical random sampling scenario on the random variable ***X***, with a probability density function (pdf) *f* (***x***; *θ*), the probability of selection is the same for each environmental combination, regardless of the value of ***x***. The pdf at ***x*** is therefore *f* (***x***; *θ*). In contrast, under a biased sampling scenario on ***X***, the probability of selecting ***x*** is proportional to a predetermined weight function *w*(***x***), implying that the pdf at the observation ***x*** is no longer *f* (***x***; *θ*). We propose a method of determining *w*(***x***) and the resulting pdf under biased sampling, which we then use to estimate a species’ fundamental niche when *f* (***x***; *θ*) is a multivariate normal density (i.e., a density based on the exponential of the negative Mahalanobis distance).

We illustrate the application of our proposed statistical model with two worked examples. In the first example, we estimate the fundamental niche of the Asian giant hornet *Vespa mandarinia*, an invasive species recorded in Europe (there is a single report in this region) and recently documented in the United States and Canada (Wilson et al., 2020). We first use presence records and an M hypothesis that only includes the native range (East and South Asia, mainland Southeast Asia, and far East Russia) of the hornet to fit a convex shape (an ellipsoid) as a model of the fundamental niche. We then evaluate the fitted model using the presence records from the invaded regions. In the second example, we again use ellipsoids, presence records, and species-specific M hypotheses to demonstrate how different M scenarios constrain a classic Mahalanobis distance model (Clark et al., 1993; Farber & Kadmon, 2003) for different hummingbird species. We identify M scenarios under which we expect the Mahalanobis distance model to deviate from the fundamental niche and instead be closer to the realized niche of a species.

## 2 Materials and Methods

### 2.1 Modeling approach

Our aim was to account for the structure of the environmental space when attempting to estimate a fundamental niche. To address this problem, we follow Austin (2002) who suggested the inclusion of three major components in any modeling procedure in ecology: (1) an ecological model that describes the ecological assumptions to be incorporated into the analysis, and the ecological theory to be tested; (2) a statistical model that includes the statistical theory and methods used; and (3) a data model that accounts for how the data were collected or measured. The ecological model, described in the following section, includes a detailed definition of the fundamental niche as a function of both fitness and the relationship between fitness and the combinations of environmental conditions in which a species has been observed. The statistical model and the data model are addressed in subsequent sections.

#### 2.1.1 The ecological model: relevant concepts in the study of the fundamental niche

The fundamental niche of a species, *N_F_*, is the set of all environmental conditions that permit the species to exist (Hutchinson, 1957; Peterson et al., 2011). Let *E* (⊆ ℝ^*d*^) be a *d*-dimensional environmental space influencing fitness (measured, for example, as the finite rate of increase in a demographic response function). Furthermore, define a function, Λ(***x***) : *E* → ℝ, that relates each environmental combination, ***x*** ∈ *E*, to fitness (Jiménez et al., 2019; Pulliam, 2000). If the fitness function has the right shape, there is a value of fitness, *λ_min_*, that can be interpreted as minimum survivorship and defines the border of the fundamental niche (Etherington & Omondiagbe, 2019; Jiménez et al., 2019). In other words, *λ_min_* is the threshold above which the fitness is high enough to support a population; any combination of environmental values that results in a fitness lower than *λ_min_* is outside the fundamental niche: *N_F_* = {***x*** ∈ *E*|Λ(***x***) ≥ *λ_min_*}. Notice that, using this notation, Λ(*μ*) = *λ_max_*.

Assumptions about the shape of a species’ response to an environmental variable (i.e., the shape of the fitness function) are central to any predictive modeling effort (Austin, 2002; Soberón & Peterson, 2019). We assume that the contour surfaces of the fitness function, Λ, and, therefore, the boundary of the fundamental niche is convex (Drake, 2015). We specifically model these level surfaces using ellipsoids in multivariate space (Brown, 1984; Jiménez et al., 2019; Maguire, 1973) because these surfaces are simple and manageable convex sets defined by (1) a vector *μ* that indicates the position of the optimal environmental conditions (the center of the ellipsoid) and (2) a covariance matrix Σ that defines the size and orientation of the ellipsoid in *E*. These two parameters can also be interpreted as the parameters of a multivariate normal density function, *f* (***x***; *μ*, Σ), in which the corresponding random variable ***X*** represents a combination environmental conditions in *E* where the species could be recorded as present (Jiménez et al., 2019). These ellipsoids are also known as Mahalanobis distance models (Farber & Kadmon, 2003) because the Mahalanobis distance defined by *μ* and Σ is equivalent to the quadratic form that defines *f* (***x***; *μ,* Σ). Ellipsoids representing the niche have been used to test the niche-center hypothesis (Osorio-Olvera et al., 2020), which predicts that the closer an environmental combination is to the vector *μ*, the higher the suitability value associated with that combination, and therefore, the higher the abundance of the species there.

We transform fitness values, Λ(***x***), by calculating 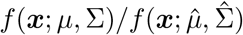 (where 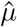 and 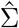 are estimates of *μ* and Σ, respectively) which produces a value between 0 and 1 that can be interpreted as a “suitability” index for the environmental combination ***x***. The central assumption here is that there is a monotonic transformation between Λ(***x***) and *f* (***x***; *μ*, Σ) in the sense that high values of fitness (near *λ_max_*) correspond to high suitability values (near *μ*). We can therefore work with the normal model to delimit the environmental requirements of a species and test the niche-center hypothesis.

Other central concepts in the study of the fundamental niche are the existing niche (Jackson & Overpeck, 2000) and the realized niche (Hutchinson, 1957), which are each subsets of the fundamental niche. The existing niche represents the existing climatic conditions in the study region, and, the realized niche represents the subset of the fundamental niche that a species occupies as a result of biotic interactions with other species. We assume that the following relationships between three niche concepts – the fundamental niche *N_F_*, the existing niche *N* *(*t*; *M*) and the realized niche *N_R_* – are fulfilled (Peterson & Soberón, 2012; Peterson et al., 2011):

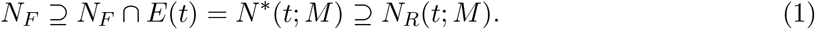

These relationships are illustrated in Figure 1, for a hypothetical species, and constitute our ecological model. In theory, every environmental combination inside the fundamental niche ellipse is suitable for the species, and, if the species can reach a site with those environmental conditions, it could persist there indefinitely (in the absence of biotic factors such as predators). However, at a given time, *t*, only a discrete subset of the environmental combinations that can be mapped into E-space exist in geographic space. This subset constitutes the *existing niche* of the species. Note that this set of environmental combinations, which are contiguous points in E-space, are divided across different regions when represented in geographic space: some of the points are in North America (purple points) and others are in South America (green points). Species have dispersal limitations that may prevent them from colonizing all the environments in their existing niche. Thus, if a species is native to North America and is only able to reach the area shaded in orange (henceforth *M*), then (i) its realized niche will be a subset of purple points (because they are suitable environments that the species has access to) and (ii) the set of green points constitutes its potential niche (because they are suitable environments that are not accessible to the species). There is therefore only a discrete set of environmental combinations from the fundamental niche that is available to the species, and we expect to observe higher abundances of the species in the environmental conditions that both exist and are close to the center of the fundamental niche ellipse.

**Figure 1:**
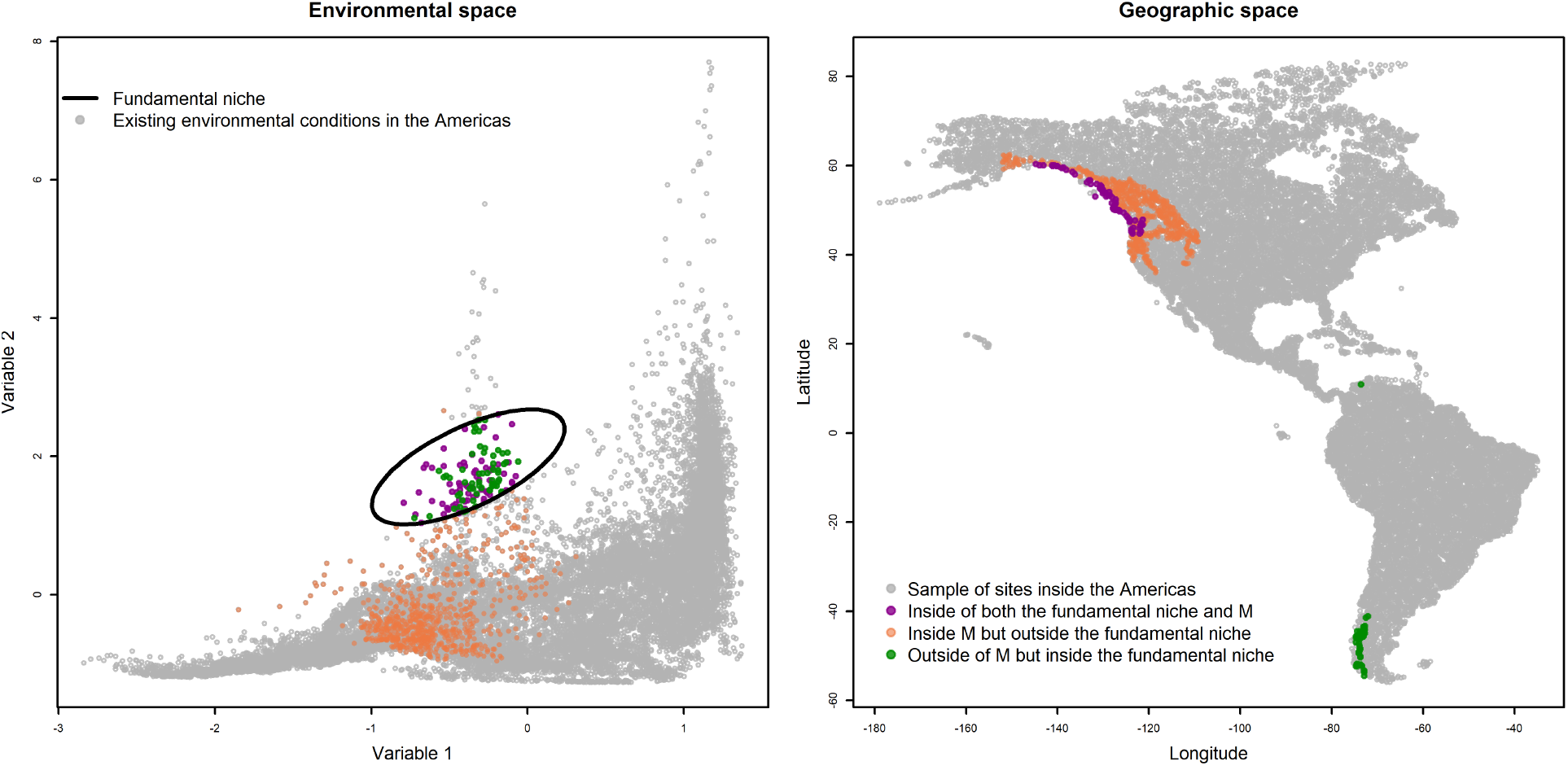
Hypothetical example of the different subsets of environmental combinations that are of interest for niche estimation. Each grey point in geographic space corresponds to a grey point in E-space, and vice versa. Left: The ellipse represents the border of a fundamental niche in which only the green and purple environmental combinations exist somewhere in geography (existing niche, *N* * (*t, M*)) and only the purple points are accessible to the species. The orange points represent environmental combinations inside M but outside the fundamental niche. Right: The corresponding regions in geographic space are highlighted with the same colors.

#### 2.1.2 The statistical and data models: two-stage sampling

Given a sample of environmental conditions where the species has been observed as present, *D* = {***x***_1_, *…, **x**_n_*}, we propose a likelihood function for the parameters that describe the fundamental niche of the species (*θ* = (*μ,* Σ)). Suppose that the environmental space *E* is defined by *d* environmental variables (i.e., *D* ⊂ *E* ⊆ ℝ^*d*^ and each point in E-space ***x**_i_* has *d* coordinates). If *E* were a uniform grid of points embedded in ℝ^*d*^, then the sample of conditions in which a species is present *D* could be considered to be a random sample of a multivariate normal variable with the density function

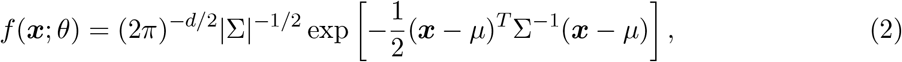

This density function could theoretically be used to define a likelihood function and estimate *μ* and Σ.

This is unfortunately not the case, for two main reasons. First, the environmental combinations that exist currently on the planet do not represent a uniform sample taken from the whole multivariate space *E* because Huthinson’s duality does not preserve distances. If we take a uniform grid in geographic space and map it into environmental space, the resulting cloud of points will be concentrated in select regions, leaving other regions empty, as seen in Figure 1. Second, species occurrences can only come from *E_M_*, or the set of environmental combinations associated with all the sites in the region that is accessible to the species *M*. The irregular shape of *E_M_* induces a sampling bias such that the probability of recording the species as present in an environmental combination is no longer given directly by *f* (***x***; *μ,* Σ), as these probabilities are affected by the availability of environmental conditions in *E_M_*.

To account for the sampling bias induced by *E_M_*, suppose that when the event {***X*** = ***x***} occurs (i.e., when the species is recorded as present at a site with these environmental conditions), the probability of observing it changes depending on the observed ***x***. We represent this probability by *w*(***x***). The set of observed environmental combinations, *D*, can therefore be considered as a random sample of the random variable ***X**_w_*, with probability density function

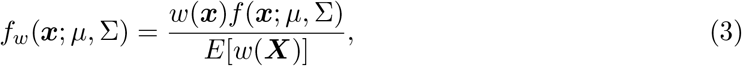

where

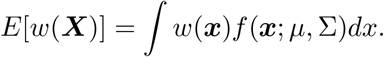

Note that *f_w_*(·) is an example of a weighted density function in which *E*[*w*(***X***)] is the normalizing factor, making the total probability function equal to unity (Lele & Keim, 2006; Patil & Ord, 1976; Patil & Rao, 1978). Patil & Rao (1978) call this normalizing factor the *visibility factor*, which captures the idea that samples from *E* are not uniform. The observed species presences can only come from *E_M_* (the visible set of environmental combinations), which may include regions in ℝ^*d*^ where the environmental combinations are abundant (associated to high values of *w*(***x***)). But, even though the points inside these abundant regions might not be close to *μ* (i.e., where *f* (***x***; *μ,* Σ) is small), the probability of observing the species in these points could be higher than the probability associated to other points closer to *μ* (i.e., where *f* (***x***; *μ,* Σ) is large) that do not exist in *E_M_* (they are not visible), or whose weight *w*(***x***) is too small.

On the other hand, equation 3 can also be described as the model resulting from a two-stage sampling design that accounts for the random process under which an environmental combination ***x*** ∈ *D* is observed (Patil & Rao, 1978). Suppose that nature produces a sample of size *N* of environmental conditions inside the fundamental niche, with probabilities of being observed given by the density function *f* (***x***; *μ,* Σ). This sample may contain any point in *E*. Because the species not only requires the right abiotic conditions to maintain a population, but also needs specific biotic conditions and can only disperse to a finite set of sites, the recorded sample will not include all the *N* observations. Instead, only a subsample of size *n < N* is selected by drawing observations from the original sample (of size *N*) with a probabilities proportional to *w*(***x***).

Without loss of generality, let us illustrate this two-stage sampling design using the fundamental niche described in the bidimensional environmental space shown in Figure 1. Suppose that a species’ fundamental niche is defined by a bivariate normal distribution with parameters:

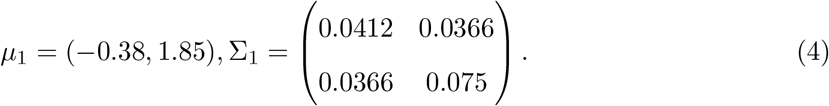

The level curves of the corresponding density function, *f* (***x***; *μ*_1_, Σ_1_), are ellipses; they are plotted in the left panel of Figure 2, where the largest ellipse corresponds to the 99% confidence region. In the first stage of the sampling process, we generate a sample of size *N* = 100 from within the fundamental niche and plot this sample on top of the ellipses. As expected, most of the environmental combinations in this sample are close to the center of the ellipses (green points inside the ellipses plotted in the left panel of Figure 2). The middle panel of Figure 2 shows all the environmental combinations that exist inside the region that is accessible to the species, *E_M_*, which are indicated by purple and orange points, in both Fig. 1 and Fig. 2). We identified the subset of points that both exist in *E_M_* and are inside the 99% confidence region of *f* (***x***; *μ,* Σ) and colored them purple. We then estimated the density function of the accessible environments, *ĥ*(·; *E_M_*), using a kernel method. The resulting level curves of *ĥ*(·; *E_M_*) correspond to the orange regions in the middle panel of Fig. 2. We used *ĥ*(·; *E_M_*) to define the weights, *w*(***x***) = *ĥ*(***x***; *E_M_*), which were used in the second stage of the sampling process. Based on the assumption that not all the *N* = 100 points from the original sample exist in *E_M_*, the second sampling stage involved selecting a subsample of size *n* = 25 from the first sample of *N* = 100 environments from the fundamental niche. The resulting sample, *D* = {***x***_1_, *…, **x**_n_*}, is shown in the right panel of Figure 2 (purple triangles).

**Figure 2:**
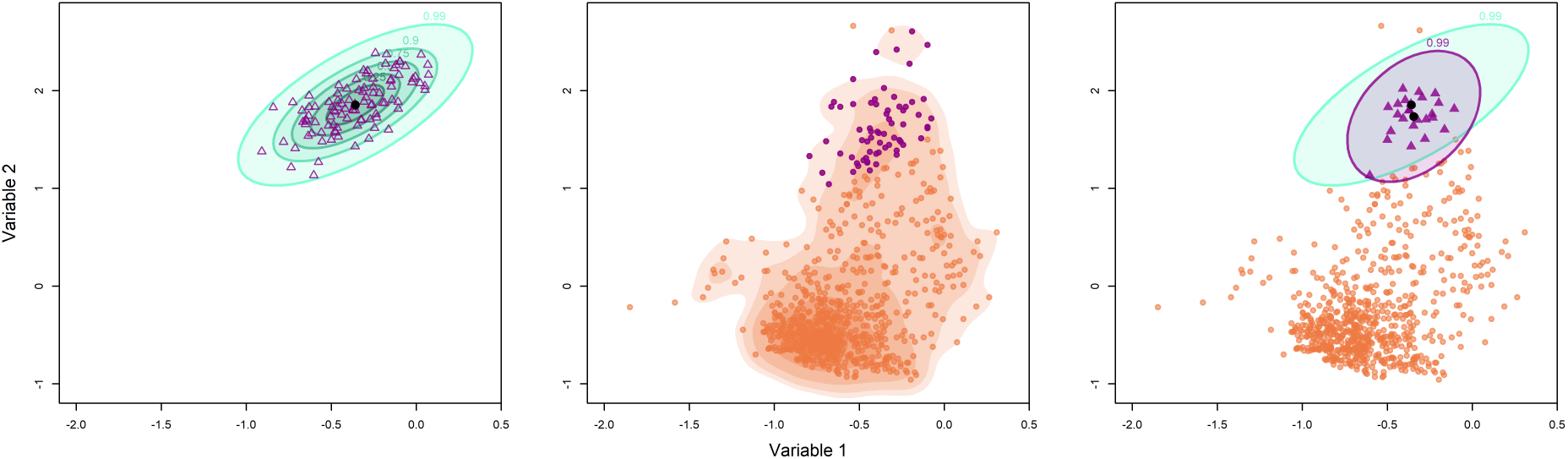
Left: A hypothetical fundamental niche represented as ellipsoids that correspond to the confidence regions of a normal distribution with parameters *μ*_1_ and Σ_1_ (see Eq.4). In the first stage of the sampling process, a sample of size *N* = 100 (purple triangles) is simulated from this distribution. Middle: The environmental combinations accessible to the species, identified as *E_M_* (orange and purple circles), and the contour levels of the kernel density function estimated using these points, *ĥ*(***x***; *E_M_*) (orange regions). Right: In the second stage of the sampling process, a subsample of size *n* = 25 (purple triangles) is selected using the weights defined by *ĥ*(=;*E_M_*). The rest of the points in *E_M_* where not in the sample (orange circles), and the hypothetical fundamental niche of the species (green ellipse) is compared against the estimated niche from a Mahalanobis distance model (purple ellipse).

Note that if we use a simple likelihood approach based on the simulated sample to estimate the parameters *μ*_1_ and Σ_1_, which is equivalent to fitting a Mahalanobis distance model, we recover a 99% confidence ellipse (violet ellipse in the right panel of Fig. 2) that is smaller than the theoretical fundamental niche of our species (largest, green ellipse in the right panel of Fig. 2). More importantly, the estimated center of this ellipse does not coincide with the optimal environmental conditions, *μ*_1_. Therefore, if we estimate the parameters that describe the fundamental niche of the species without accounting for the distribution of available environments, we cannot claim that the model fully recovers the species’ fundamental niche. Moreover, if we attempt to test the niche-center hypothesis using an ellipse recovered from a Mahalanobis distance model, we will expect high abundances around an environmental combination that is not the true optimum.

With respect to the application of the two-stage sampling method, we assume that the sample of environmental combinations where the species of interest was observed as present, *D*, is a random sample of the random variable ***X**_w_* with a probability density function given by Eq. 3. We can therefore define the likelihood function of the parameters of interest *θ* = (*μ,* Σ) as follows:

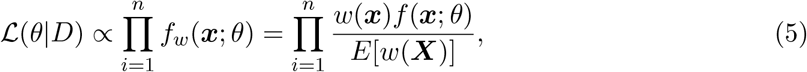

where the function *w*(***x***) is approximated using all the environmental combinations accessible to the species along with a kernel density procedure, and we obtain a Monte Carlo estimate for the expected value *E*[*w*(***X***)] (which was defined as an integral; see Eq. 3). Note that the function *w*(***x***) does not depend on the parameters of interest and can be ignored when maximizing the log-likelihood function. Furthermore, because the analytical form of *w*(***x***) is unknown, the analytical form of *E*[*w*(***X***)] is also unknown. However, we can sample environmental combinations from the availability distribution by randomly choosing sites inside M and extracting the values of the environmental variables at those sampled sites. Using this sample, we can obtain Monte-Carlo estimates of *E*[*w*(***X***)] for any fixed value of *θ*. This method has been used before in the statistical modeling literature and is called the method of simulated maximum likelihood (Lele & Keim, 2006; Robert & Casella, 1999). We can use this method to obtain a Monte Carlo estimate of the log-likelihood function as follows:

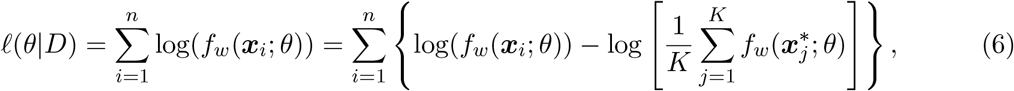

where 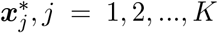 is a random sample that was sampled with replacement from the distribution *w*(***x***). The size of this sample, *K*, must be large enough to ignore Monte Carlo error. Once this sample is generated, we can apply standard optimization techniques to minimize the negative log-likelihood function in Eq. 6 and obtain the maximum likelihood estimators 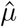 and 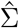.

In summary, our proposed modeling procedure uses a sample of occurrences for the species of interest together with polygons that represent the geographic areas accessible to the species. Inside these polygons, we extract environmental values from sites chosen at random. This is done with two different purposes: (1) to estimate a kernel density used to define the weights in the likelihood function, and (2) to obtain a Monte Carlo estimate of the log-likelihood function. Once we have all the elements of the likelihood function, we calculate the maximum likelihood estimates of the parameters 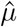 and 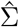. These estimated parameters will allow us to plot ellipses in E-space that represent the border of the estimated fundamental niche. In all the examples we show, we plot the ellipses that correspond to the 99% confidence regions of the fitted multivariate normal distribution. We compare these ellipses to the ones that correspond to the 99% confidence region of a standard Mahalanobis distance model (Farber & Kadmon, 2003) with estimated parameters 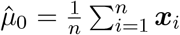 and 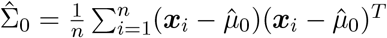 (i.e., the maximum likelihood estimates under a multivariate normal model described in Eq. 2). We hypothesized that the two ellipses will be similar in cases where *E_M_* covers most of the fundamental niche of the species (i.e., the overlap between these two sets is high) and the distribution of points in *E_M_* is approximately uniform.

Finally, we project the resulting models back to geographic space. Once we have the maximum likelihood estimates of the parameters of interest, 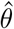, we use the multivariate normal density function given in Eq. 2 to calculate a suitability index, that can be plotted in G-space as either a continuous value or a binary region defined by a threshold. For interpretation purposes, it is convenient to standardize this suitability index to the interval (0, 1), which is easily done by dividing *f* (***x***; *θ*) by its maximum value, 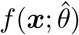.

### 2.2 Data

Using our method to approximate fundamental niches requires three types of data: (*i*) species occurrence data, which should go through a standard process of detecting and removing inaccurate records before being used (see Cobos et al. (2018) for an example of this); (*ii*) an M hypothesis or geographic polygon that encloses all the sites accessible to the species, taking into consideration the dispersal ability of the species and natural geographic barriers; and (*iii*) environmental layers, cropped to the study area, from which we can extract environmental data for the occurrences and for the sites inside M. In the following sections, we describe the datasets that we used to create our worked examples.

#### 2.2.1 Occurrence data

We selected seven species to illustrate the use of the statistical model that we present here. The first species is the Asian giant hornet *Vespa mandarinia* (Matsuura & Sakagami, 1973; Matsuura, 1988). Because *V. mandarinia* is an invasive species, using it in our model can provide valuable insight into whether the niche estimated by our model is a good approximation of a true fundamental niche. In other words, we test whether the estimated fundamental niche contains the locations that this species has been able to invade. We also selected six species of hummingbirds for a separate analysis: *Amazilia chionogaster*, *Threnetes ruckeri*, *Sephanoides sephanoides*, *Basilinna leucotis*, *Colibri thalassinus*, and *Calypte costae*. These species were used because detailed M hypotheses are available for them.

The occurrence data for *V. mandarinia* was obtained from the Global Biodiversity Information Facility (GBIF) database (https://www.gbif.org/). We downloaded 1944 occurrence records for (GBIF, 2020), which, after undergoing a standard cleaning procedure (Cobos et al., 2018), were reduced to 170 presence records in the species’ native range and one presence record in Europe. To avoid introducing an extra source of bias (Anderson, 2012), the cleaned dataset was spatially thinned by geographic distance (at least 50 km away), resulting in a final sample of 46 presence records that were used to fit the niche models (see red points in Fig. 3).

**Figure 3:**
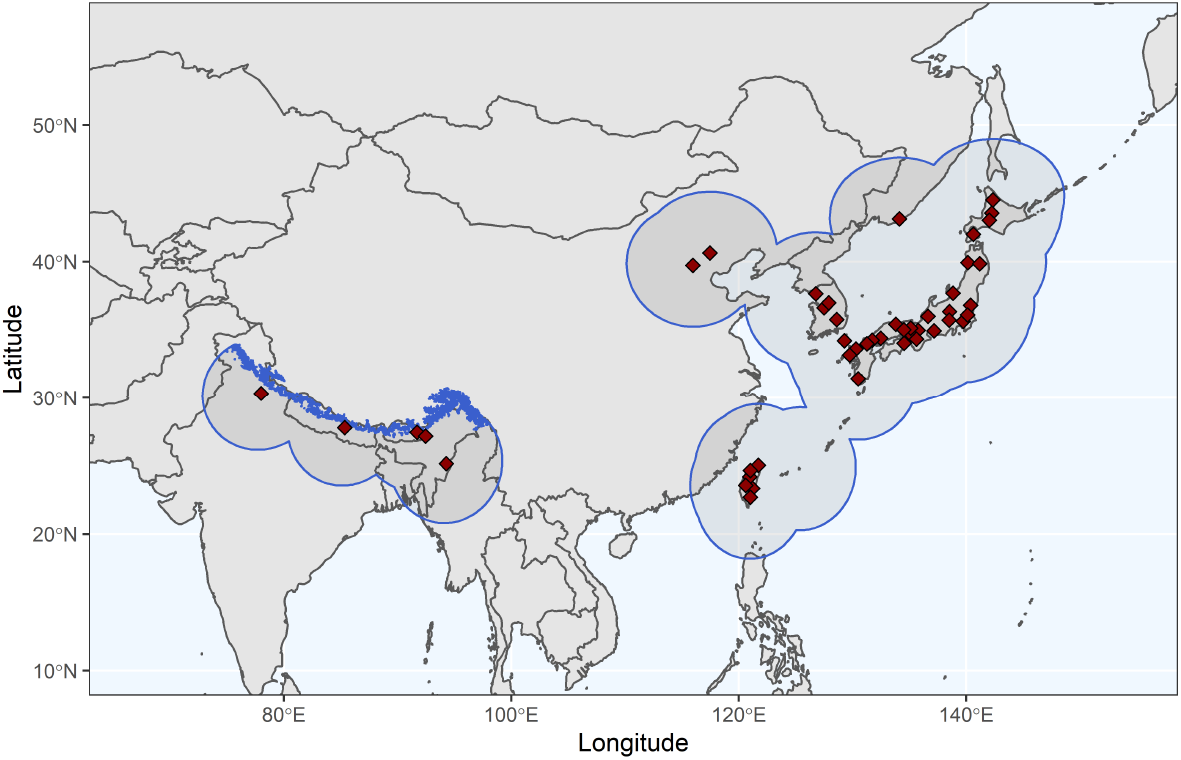
Occurrence records (red squares) of *V. mandarinia* in its native range, and the M hypothesis (regions delineated with blue lines) created from a combination of buffers and the known elevation range of the species. There were a total of 46 occurrence records after the data cleaning and thinning process.

For the hummingbird species, we used the cleaned occurrence records that were used by Cooper & Soberón (2018), which are available at https://github.com/jacobccooper/trochilidae and whose original source is also GBIF. Cooper & Soberón thoroughly revised and cleaned these presence records, and eliminated misidentified individuals and synonyms. They additionally defined the regions that were accessible to most of the extant hummingbird species and used those regions to fit species distribution models. Given the success of Cooper & Soberón at using these M hypothesis, we believed their data provided a great opportunity to test our proposed model. Figure 4 shows the presence data of each hummingbird species. The sample sizes range from 148 observations for *A. chionogaster*, to 926 presences for *C. costae*. We selected these six species because they occupy different regions of the Americas, from the southern United States to the Patagonia. Given this wide geographic range, we believed it would be interesting to compare the estimated niches of all the species in E-space and evaluate how similar or different the optimal environmental conditions are among the different species. Although the different species are likely to occupy different regions of E-space, we suspected that their fundamental niches might share some environmental combinations (niche overlap).

**Figure 4:**
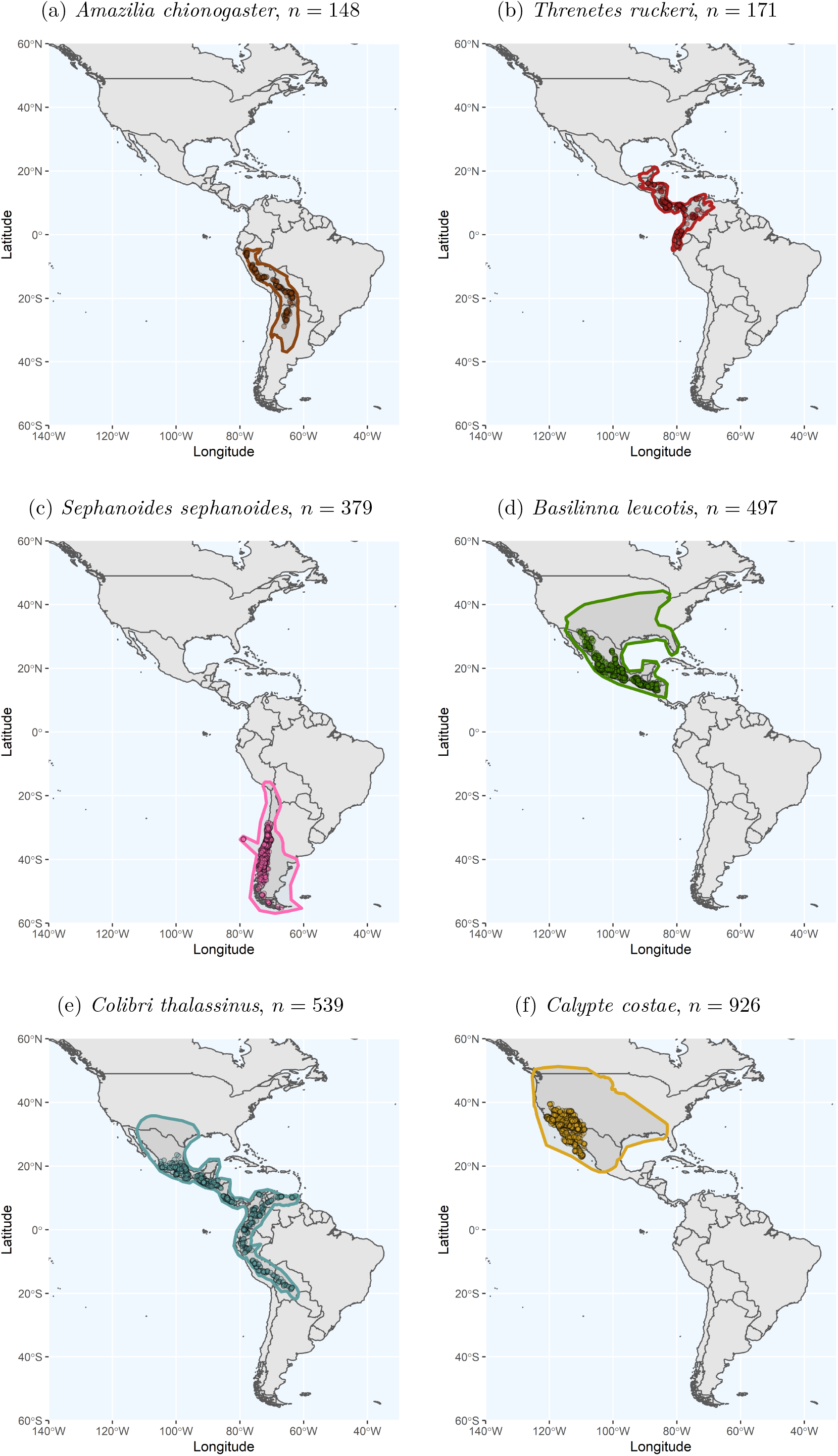
Occurrence samples (dots) and M polygons (estimated accessible areas around the occurrences)for the six species of hummingbirds: (a) *A. chionogaster* in brown, (b) *T. ruckeri* in red, (c) *S. sephanoides* in pink, (d) *B. leucotis* in green, (e) *C. thalassinus* in pale blue, and (g) *C. costae* in yellow. The sample sizes (*n*) are given in each panel.

#### 2.2.2 M hypothesis

In the case of *V. mandarinia*, we defined the area accessible to the species as a combination of buffers that represent the species’ dispersal ability and the elevation range where the species is known to occur (850 - 1900 m). First, we identified the region delineated by a buffer of 500 km around all occurrence records in the sample, which accounts for the dispersal ability of the hornets (Matsuura & Sakagami, 1973; Matsuura, 1988). Second, we clipped this region with an elevation layer to remove regions at elevations higher than 1900 m. The resulting polygon is outlined in blue in Figure 3. Note that this polygon includes some sites not on the continent; however, we only extracted the environmental values for inland sites within this blue M, and did not considered oceanic regions.

For the hummingbird species, we used the polygons generated by Cooper & Soberón (2018). Polygons were hypothesized based on the known species occurrences, topography, ecoregions, and estimated dispersal distances, and where bounded by significant geographical barriers such as large rivers and mountains. By doing so, they took into account all the criteria that are know to yield more accurate models (Barve et al., 2011; Owens et al., 2013; Owens et al., 2020; Saupe et al., 2012). Figure 4 shows the six polygons for the different species. Note that some species, like *T. ruckeri*, occupy most of their accessible area (Fig. 4b), whereas others, like *B. leucotis* and *C. costae*, occupy only a fraction of it. In addition, there is wide diversity in range sizes, with *S. sephanoides* having a more restricted range compared to *C. thalassinus*. Given this geographic variation among species, we were interested in making similar comparisons in E-space.

#### 2.2.3 Environmental variables

The climatic layers used to create the models for each species came from the WorldClim database (Hijmans et al., 2005), which provides open access to bioclimatic layers built with monthly average measurements for the period 1970-2020. We used only two of the 19 variables available in this database: annual mean temperature (Bio1), and annual precipitation (Bio12). Both variables were recorded at 10 arcmin resolution. We clipped each climatic layer using the polygons that correspond to the M hypotheses for each species.

These two climatic variables variables were selected because they are known to be biologically meaningful for both the Asian hornet and the selected hummingbird species (Root, 1988). Specifically, it has being documented that the wasps of the genus *Vespula* have high endothermic capacity which benefits their thermoregulatory efficiency allowing them, at the same time, to survive to broad temperature ranges (Käfer et al., 2012). Furthermore, other studies suggest that the queens of this genus prefer green environments, such as agricultural and forested areas, or parks (Kim et al., 2020); whose presence is highly correlated with the amount of rainfall in the area. Furthermore, in the case of the hummingbirds, we wanted to compare their estimated niches. Because comparisons of estimated niches in E-spaces with different axes and/or dimensions are not possible, we only looked at these two dimensions of their niches.

All analyses were performed in R version 3.6.3 (R Core Team, 2020). We used several existing packages for data preparation, analysis, and visualization: ggplot2 (Wickham, 2016), ks (Duong, 2020), raster (Hijmans, 2020), rgdal (Bivand et al., 2020),rgeos (Bivand & Rundel, 2019), scales (Wickham & Seidel, 2020), sf (Pebesma, 2018). We additionally created new functions that can be used to reproduce our examples, and apply our methodology to other species. These functions can be found at https://github.com/LauraJim/Nf_model_with_M.

## 3 Results

### 3.1 Estimated fundamental niche of *Vespa mandarinia*

We randomly sampled 10,000 sites inside the accessible (native) region of the Asian giant hornet (shown in Fig. 3) and extracted their environmental values (*E_M_*) so we could plot them in E-space. The resulting set *E_M_* is shown in Figure 5 as a cloud of points in the background (grey open circles). We used this *E_M_* and the 46 presence records of *V. mandarinia* (red points in Fig. 5) to determine the log-likelihood function (Eq. 6) of the parameters that describes its fundamental niche under the weighted-normal model, *θ* = (*μ,* Σ). We maximized this log-likelihood function to obtain the maximum likelihood estimates (MLEs) 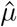 and 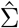. For comparison, we also used the 46 presence records only to obtain MLEs using the Mahalanobis distance model, 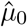 and 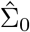. The resulting MLEs for both models are given in Table 1 and the corresponding 99% confidence regions are plotted in Figure 5.

**Figure 5:**
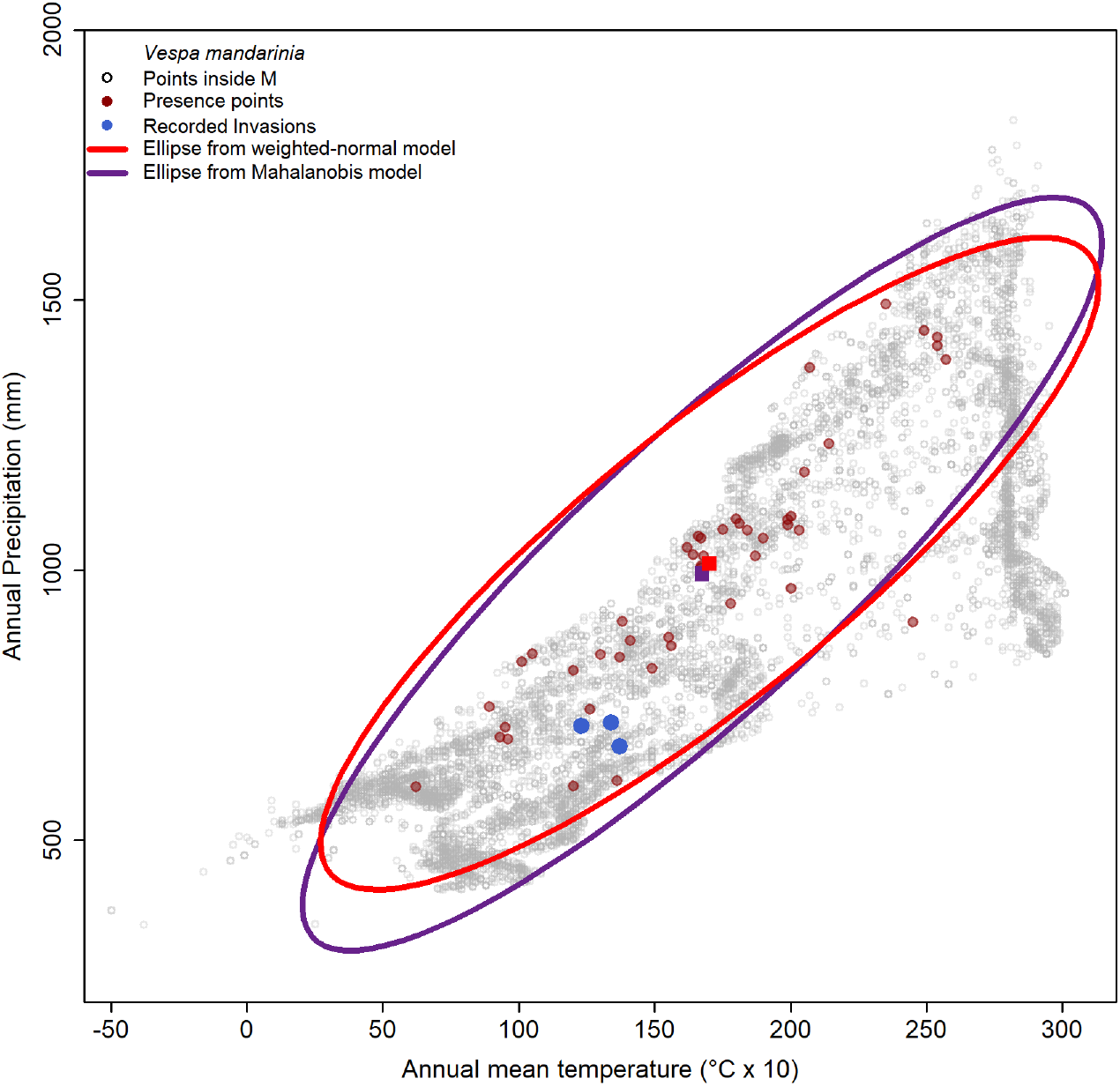
Estimated ellipses (99% confidence regions) from the weighted-normal model(red) and the Mahalanobis distance model (purple), which are defined by 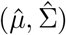 and 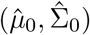, respectively, as given in Table 1. Red points are the presences from the native range of *V. mandarinia* that were used to fit the models, and blue points are presences recorded outside the native range. The centers of the ellipses are indicated with a square of the same color as the corresponding model. The grey points represent *E_M_*.

**Table 1:** MLEs of the weighted-normal model (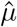 and 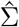) and the Mahalanobis distance model (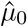 and 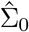) obtained from 46 presences of *V. mandarinia* inside its native range.

| Model | Weighted-Normal |  | Mahalanobis |  |
| --- | --- | --- | --- | --- |
| Parameters | $\hat{\mu}$ | $\hat{\Sigma}$ | $\hat{\mu}_0$ | $\hat{\Sigma}_0$ |
| *V. mandarinia* | (170.21, 1012.19) | $\begin{pmatrix} 2223.45 & 8031.54 \\ 8031.54 & 39554.33 \end{pmatrix}$ | (167.52, 992.97) | $\begin{pmatrix} 2348.47 & 9813.12 \\ 9813.12 & 52770.37 \end{pmatrix}$ |

The MLEs of *μ* (the optimal environmental conditions) estimated by both the weighted-normal model and the Mahalanobis distance model, are highly similar (purple and red squares in Figure 5). However, the models predicted different limits for the fundamental niche of *V. mandarinia*. Specifically, the estimated *N_F_* is larger under the Mahalanobis distance model (purple ellipse in Fig. 5), whereas the variance predicted by the weighted-normal model for annual precipitation is smaller, as shown in Table 1. Nevertheless, both estimated ellipses contain 45 out of the 46 initial presence points and they contain all the records from the locations outside the native range of *V. mandarinia* (blue points in Fig. 5). The presences from the invaded regions are not close to the center of the ellipses, but they are placed in a region of *E_M_* where the environmental combinations are well represented in G-space.

We calculated a standardized suitability index for the Asian giant hornet using the MLEs from the weighted-normal model and Eq. 2. Using this index, we created a worldwide suitability map (Fig. 6) to visually assess if there are regions where *V. mandarinia* could theoretically establish based solely on the species’ environmental requirements. As shown in Figure 6, the Pacific coast of the northern United States and southern Canada, where the species has already been confirmed, has a low to moderate suitability index. Similarly, most of Europe has a low to moderate suitability, particularly around the one site where the species was already recorded. Additional maps that are focused more closely on the West Coast of North America, Europe, and the native range of *V. mandarinia* are provided in the Supplementary Material. We also note that there are other regions within the Americas that appear highly suitable for *V. mandarinia*, such as the eastern United States, the mountain ranges of Mexico and the Andes, and the Brazilian and Ethiopian highlands.

**Figure 6:**
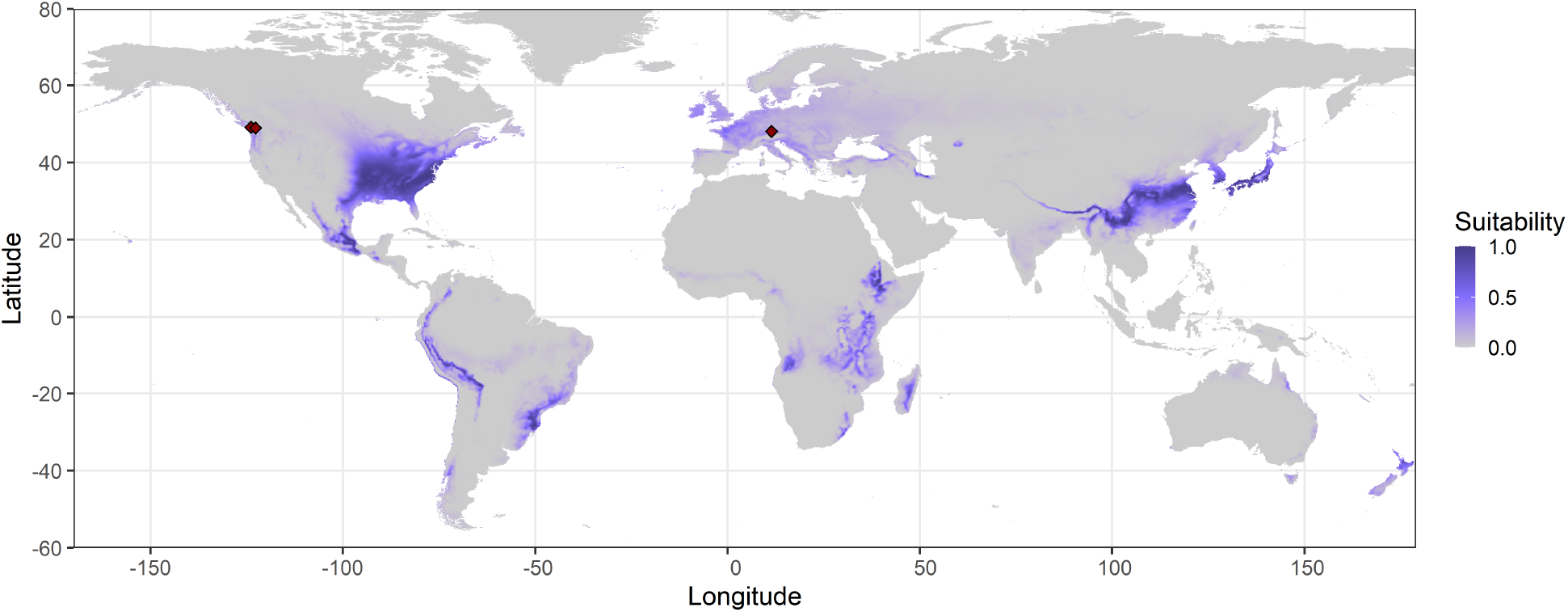
Worldwide suitability map of *V. mandarinia*. Dark purple regions correspond to highly suitable sites where the environmental combinations in E-space are close to the center of the estimated fundamental niche, and light purple regions correspond to sites with low suitability whose environmental combinations in E-space are near the border of the fundamental niche. The red dots indicate sites outside the species native range where the species has been reported.

### 3.2 Estimated fundamental niches of hummingbird species

In the case of the six hummingbird species that we chose for our analysis, the accessible areas (Ms) and the presence records were in different regions of G-space but also, of E-space. The corresponding *E_M_* and presence points therefore occupied different regions (see grey clouds of points in Fig. 7). Using the M hypothesis and the presences, we estimated the parameters *μ* and Σ, which describe the fundamental niche of each species, using both the Mahalanobis distance model and the weighted-normal model. The resulting MLEs for each species are given in Table 2.

**Table 2:**
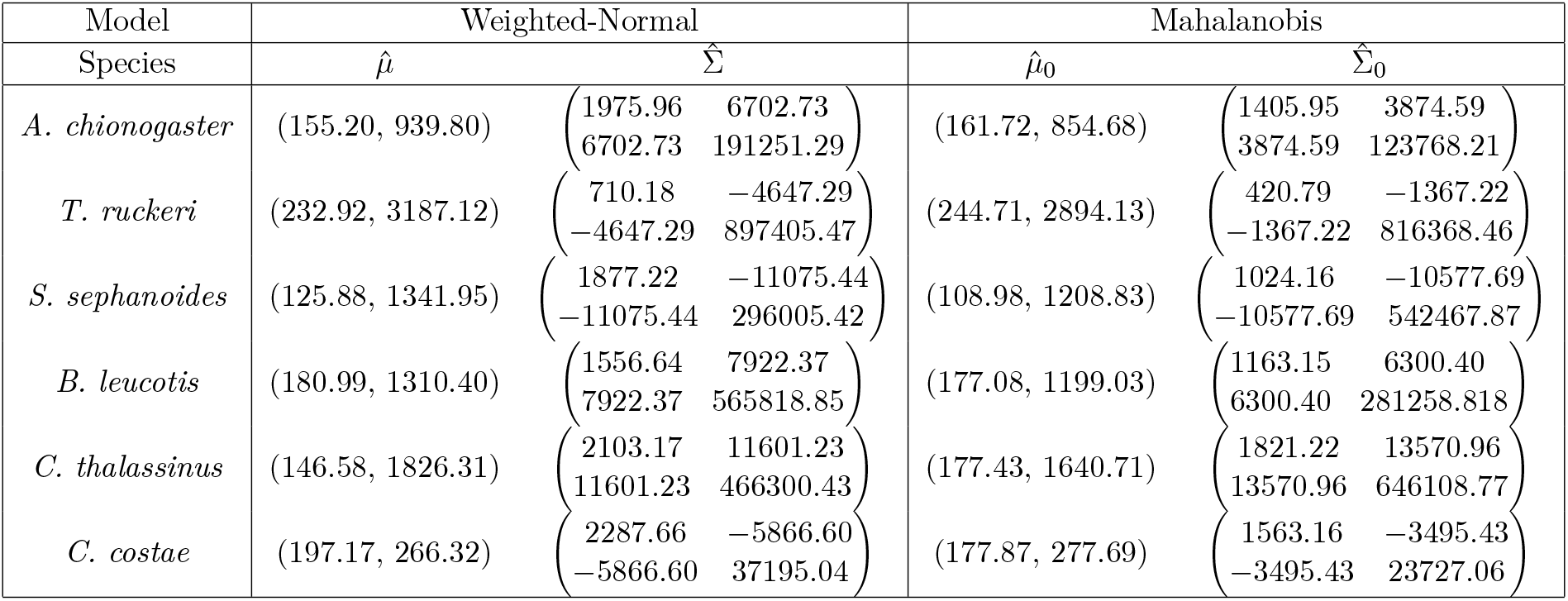
Maximum likelihood estimates of the parameters that determine the fundamental niche of the six hummingbird species. Parameters were obtained with the weighted-normal model (second column) and the Mahalanobis distance model (third column).

**Figure 7:**
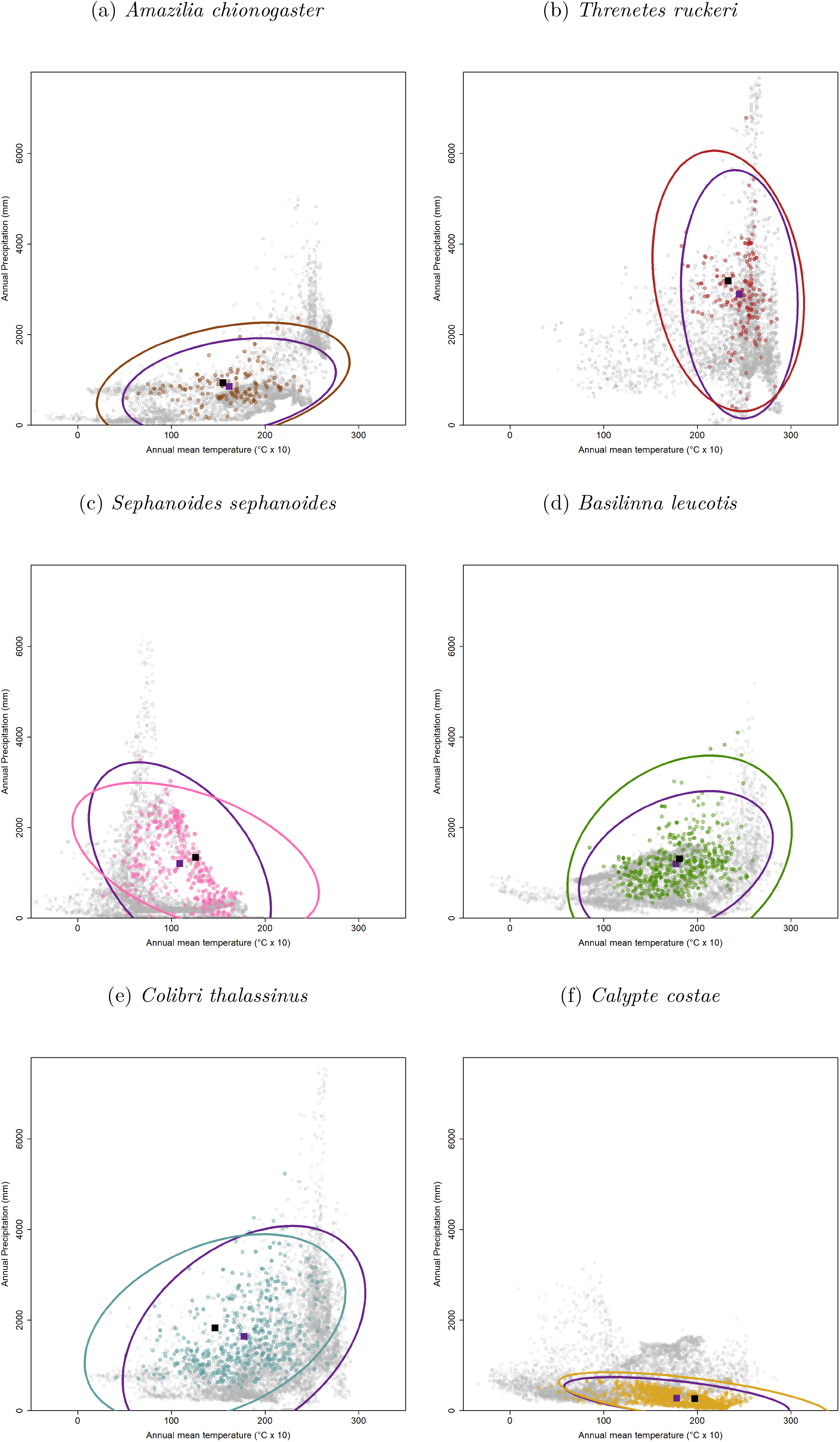
Estimated fundamental niches for the six species of hummingbirds. In all panels, the purple ellipse represents the estimated niche from a Mahalanobis distance model, and the other ellipse represent the estimated niche from our proposed weighted-normal model. The centers of both ellipses are marked with a purple square and a black square, respectively.

Unlike the *V. mandarinia* example, the weighted-normal model predicted broader fundamental niches for most of the studied hummingbirds except for *C. thalassinus*. The estimated centers of the ellipses (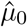 and 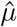) and their orientation with respect to the axes were similar for *A. chionogaster* and *B. leucotis*, however, there was a clear difference between the centers of the estimated ellipses for the *T. ruckeri*, *S. sephanoides*, *C. thalassinus*, and *C. costae*.

For all the hummingbird species, both the Mahalanobis distance and the weighed-normal models agreed on the sign of the covariance between the two environmental variables used to describe and compare the fundamental niches. The estimated ellipses for *S. sephanoides* were the most different with respect to the magnitude of the estimated covariance between the two environmental variables. We additionally noted that, with the exception of *C. thalassinus*, the ellipses obtained with the weighted-normal model contained more presence points than the ellipses obtained from a simple Mahalanobis distance model.

It is also worth noting that, under the Mahalanobis distance model, the estimated optimum temperature for four of the six hummingbird species was between 16 and 18 degrees Celsius (see Table 2). The Mahalanobis distance model thus predicts that the species temperature optima are not substantially different among these species, even though they differ in their precipitation optima. However, under the weighted-normal model, the estimated optimum temperatures were clearly different among all the species (see Figure 8).

**Figure 8:**
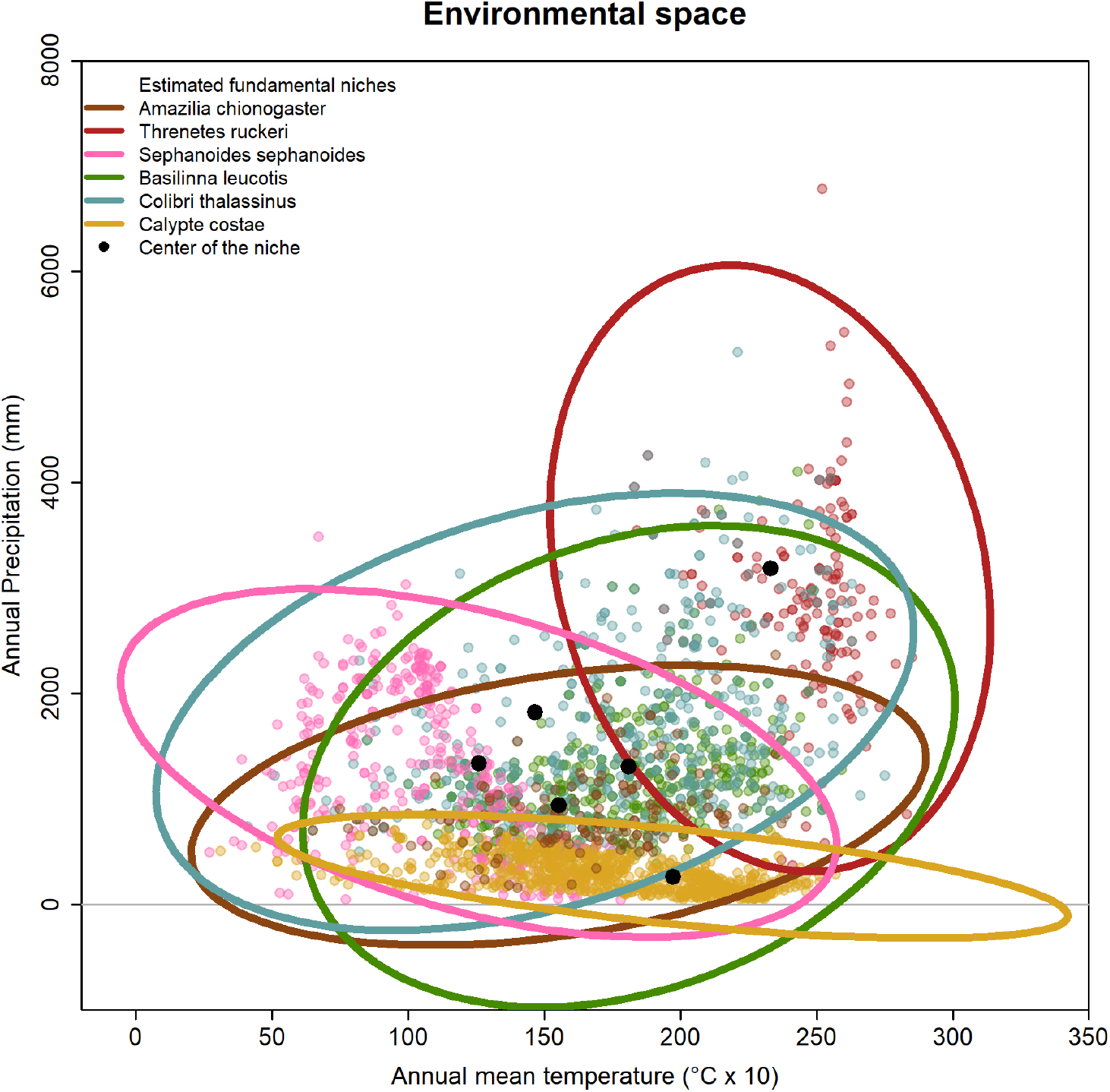
Comparison of the estimated fundamental niches (ellipses) for the six species of hummingbirds and the presence data used to fit the models. The centers of the ellipses are marked with a black circle

## 4 Discussion

The calibration region limits the presence of a species to a discrete set of environmental combinations, and the probabilities of sampling different environments from this set are not uniform. Using kernel methods, we modelled this non-uniform E-space and used the kernel as a weight function in a multivariate normal model, which we then applied to estimating the fundamental niche of a species of interest. Our main result is that accounting for the shape of the E-space defined by an *M* indeed affects the estimation of the ellipsoids we use to model fundamental niches.

We illustrated the application of our proposed method with, first, the Asian giant hornet (*Vespa mandarinia*, invasive to North America). The ellipse fitted with our model, which represents the border of the fundamental niche of *V. mandarinia*, contained the sites outside the species native range where the species has being reported (see red dots in Fig. 5). Moreover, our results agree with other modeling exercises in predicting a low to moderate suitability for the Pacific Northwest coast of the United States and Canada, and over Europe (Alaniz et al., 2020; Moo-Llanes, 2021; Norderud et al., 2021; Nuñez-Penichet et al., 2021; Zhu et al., 2020), as well as, on identifying a significant proportion of the East coast of the United States as an area of high concern (Fig.6).

Studies that have adopted Maxent as their modeling framework often have to fit a large set of candidate models (using different combinations of variables, resampling techniques, and parametrizations) and apply different selection criteria to generate a final model; all this to counteract for Maxent’s known tendency to fit arbitrarily complex models (Warren & Seifert, 2011). For example, Nuñez-Penichet et al. (2021) fitted 4,560 Maxent models using raw variables and 880 candidate models using PCs; while Moo-Llanes (2021) fitted candidate models using four different strategies of variable selection combined with 17 possible values for the regularization parameters, and 29 possible combinations of five feature classes from Maxent. Unlike these studies, our modeling framework relied on having two biologically significant environmental variables for the species and a reasonable approximation of its accessible area to arrive to very similar conclusions, and no parameter or variable tuning was needed beforehand.

In the second example, our estimates of fundamental niches were used to test the hypothesis of niche conservatism among closely related species (which is about the fundamental niche and predicts low niche differentiation between species over evolutionary time scales (Peterson et al., 1999)). We examined six hummingbird species that according to phylogenetic taxonomies belong to different major clades: *T. ruckeri* in the Hermits (Phaethornithinae), *C. thalassinus* in the Mangoes (Polytmini), *S. sephanoides* in the Coquettes (Lophornithini), *C. costae* in the Bees (Mellisugini), *A. chianogaster* and *H. leucotis* in the Emeralds (Trochilini). The clades here were listed starting with the one having the oldest split and following the order in which the splitting continued (Hernández-Baños et al., 2014; McGuire et al., 2009). We compared the estimated fundamental niches (ellipses) of these six species with respect to the phylogenetic relationships among them and noted that *T. ruckeri*, which has the oldest split in the phylogeny, had the most different estimated fundamental niche (red ellipse in Fig. 8) among the six species. On the other hand, the other five species have fundamental niches that are more similar, their optimal environmental conditions are close and their elliptical borders intersect significantly.

### 4.1 Future directions

Given that we use the distribution of *E_M_* to inform the model about the uneven availability of environmental combinations induced by M, the delimitation of M is crucial. However, there is no unique way to delimit the accessible area of a species (Barve et al., 2011). When outlining M, we need to simultaneously consider several factors: the natural history of the species, its dispersal characteristics, the geography of the landscape, the time span relevant to the species’ presence, and any environmental changes that occurred during that time span, among other things. Although we do not favor any particular approach as the best method for estimating M, we acknowledge that all those factors have an important role in determining its shape. As a future work of this research project, a thorough analysis comparing different approaches to outlining M and testing their effects in recovering the fundamental niche of a species under the proposed model is needed.

The presence of geographic sampling biases in primary biodiversity data and the implications of these biases (e.g., decreasing model performance) are broadly recognized (Kadmon et al., 2004; Meyer et al., 2016). Here, we focused on one specific type of bias that affects the estimation of a fundamental niche. However, other types of sampling biases, such as the accessibility of a site to observers, may also affect the estimation of the fundamental niche. In such cases, we would still use a multivariate normal distribution to represent the hypothesis that the fundamental niche has a simple, convex shape, but we would modify the way we define the weights in the likelihood function (Eq. 5). For example, suppose we want to include the effects of two sources of bias that affect the presence of a species: the bias induced by M and the differences in sampling intensity across the landscape due to differences in human accessibility (i.e., accessibility bias). If the sources of bias can be considered as independent, we can calculate a single, general sampling probability distribution, to define *w*(***x***), as the product of two probabilities: the probability of observing ***x*** considering the distribution of points in *E_M_* and the probability of observing ***x*** considering the accessibility bias. We proposed using a kernel to estimate the former, and Zizka et al. (2020) recently developed a method to quantify accessibility biases that could be used to determine the later.

Several other generalizations of our method are possible. For example, information from physiological experiments could be included if we consider the Bayesian approach proposed by Jiménez et al. (2019). To do so, we need to transform the likelihood function (Eq. 5) into a posterior distribution that uses the tolerance ranges obtained from physiological experiments to define the *a priori* distributions of the parameters *μ* and Σ (*g*_1_(*μ*) and *g*_2_(Σ), respectively), as follows:

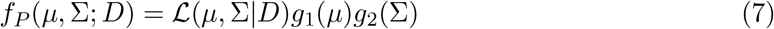

where

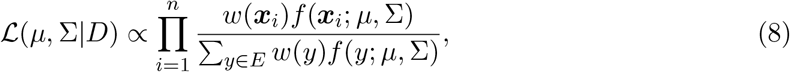

and the function *f* (·; *μ,* Σ) is given by Eq. 3. Jiménez et al. (2019) also showed how to modify *f* (·; *μ,* Σ) to get asymmetrical confidence regions when species that are known to have asymmetrical response curves for the environmental variables under consideration. Furthermore, both the likelihood and the Bayesian approaches can be applied in cases where it makes sense to include more than two environmental variables in the delimitation of a species’ fundamental niche. The disadvantage of making these modifications to our proposed model is that they increase the algebraic complexity of the model and the number of parameters to be estimated, and more computational power is therefore needed to either maximize the likelihood function or simulate from the posterior distribution.

## 5 Conclusions

In recent decades, we have seen a significant increase in the availability of species presence data and software for ecological niche modeling and species distribution modeling. At the same time, however, we have seen a concomitant increase in the number of studies that question important conceptual and methodological aspects of ENMs (Austin, 2002; Godsoe, 2010; Jiménez-valverde et al., 2008; Lobo, 2008). For that reason, we agree with Jiménez-valverde et al. (2008), who concluded that the lack of a solid conceptual background endangers the advancement of the field. The development and application of ENM/SDM should be rooted in a firm understanding of the technique’s conceptual background. We intended to lead by example in this work by explicitly relating the ecological theory to the statistical method that we developed to estimate the fundamental niche of a species.

It is universal practice in SDM and ENM to use the environmental combinations observed at the occurrence sites to fit models. If our objective is to use presence data to approximate the fundamental niche, it is essential to acknowledge that species’ presence data come from the realized niche which implies that additional information is needed to determine the whole environmental potential of the species (Qiao et al., 2018). In this study, we emphasized that we must account for the sampling bias induced by the available E-space in the region of interest. We have therefore proposed defining the distribution of environments that are accessible to the species based on the dispersal ability of the species and geographic barriers to dispersal. This is a definition of *M*, the geographic area accessible to the species via dispersal (*sensu* Peterson et al., 2011), but here we highlighted the implications of sampling from M in E-space, which are seldom discussed.

## Acknowledgements

We thank the KU-ENM group at the Biodiversity Institute of the University of Kansas for providing valuable comments. We would also want to thank Maria E. Orive at the University of Kansas for her valuable comments. L.J. was supported by the Mexican Council of Science and Technology (CONACyT) grant 409052.

## Supplementary materials

In the main text, we provided a worldwide suitability map for *V. mandarinia* obtained using the weighted-normal model that we have proposed for estimating the fundamental niche of a species. Here, we provide three additional maps focused on areas of concern for this invasive species: (1) the native range of the species in Figure S1; (2) Europe in Figure S2; and (3) the West Coast of North America in Figure S3. The same color scale used in Figure 6 is used in these figures: dark purple correspond to highly suitable sites where the environmental combinations in E-space are close to the center of the estimated fundamental niche, and light purple corresponds to sites with low suitability where the environmental combinations in E-space are near the border of the fundamental niche.

**Figure S1:**
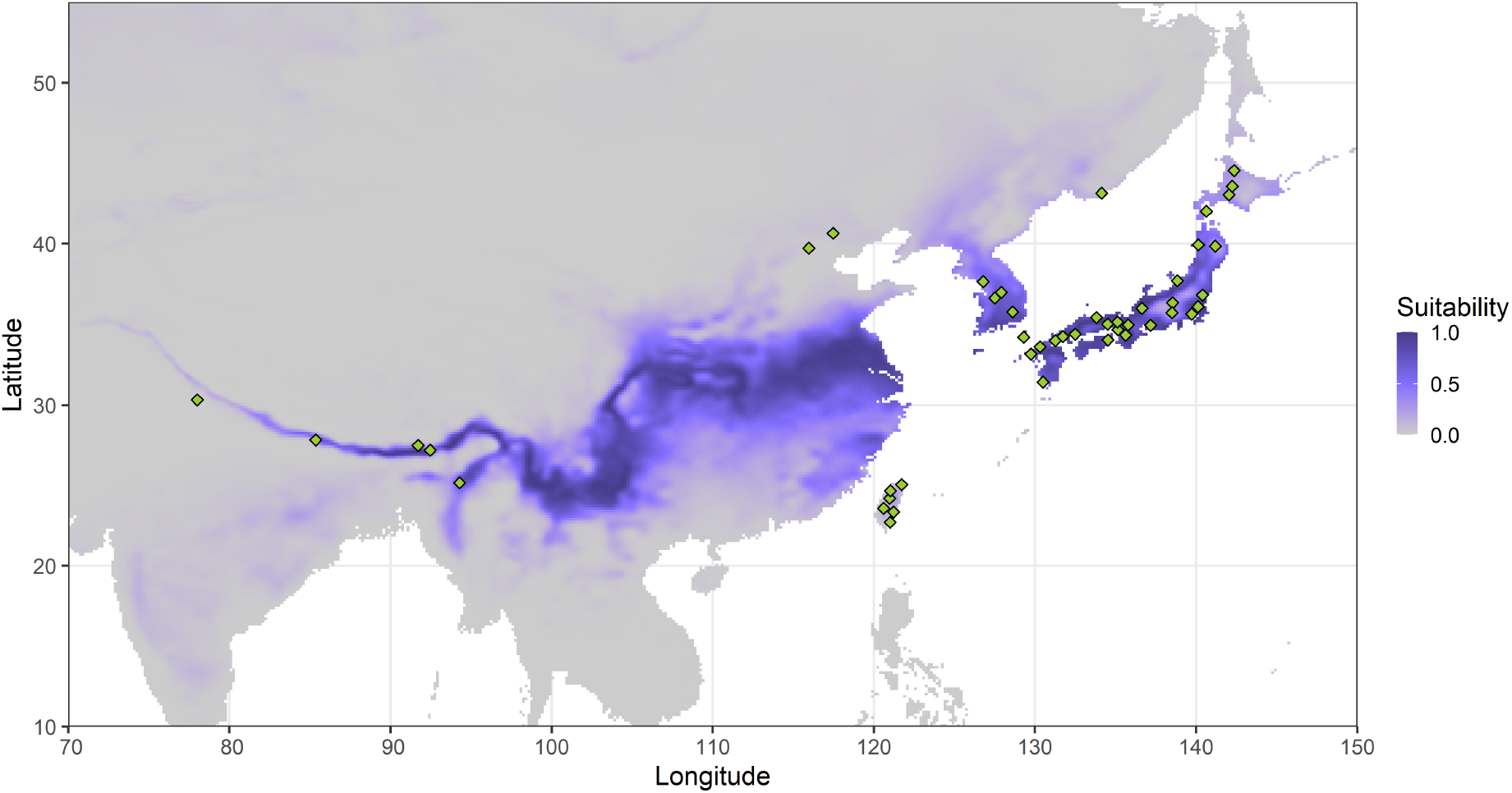
Suitability index obtained with the weighted-normal model plotted in the native range of *V. mandarinia*. The green squares are the occurrence points used to estimate the fundamental niche of the species.

**Figure S2:**
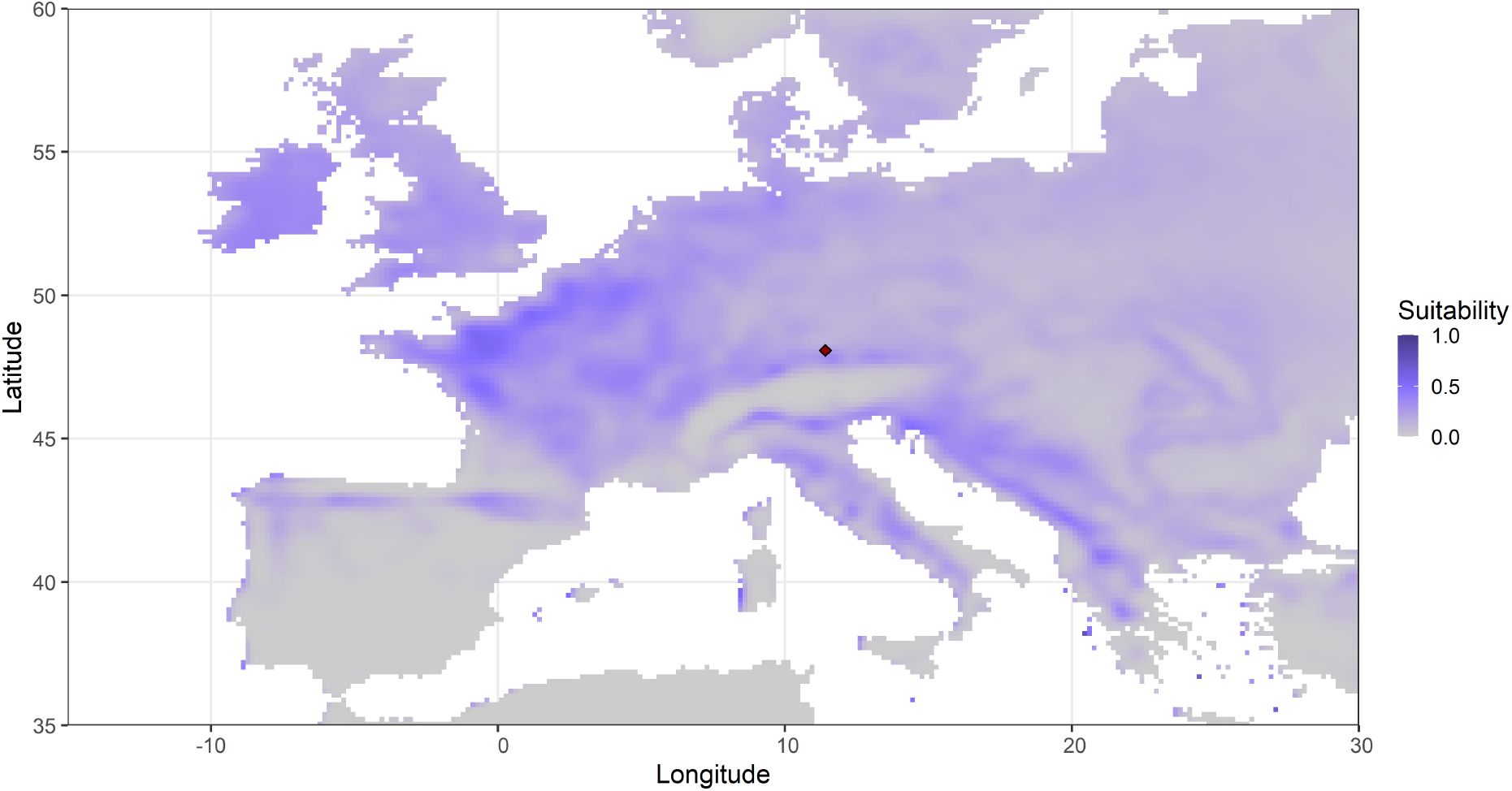
Suitability index obtained with the weighted-normal model for *V. mandarinia* plotted in Europe. The red square is an observation of the species recorded in Germany.

**Figure S3:**
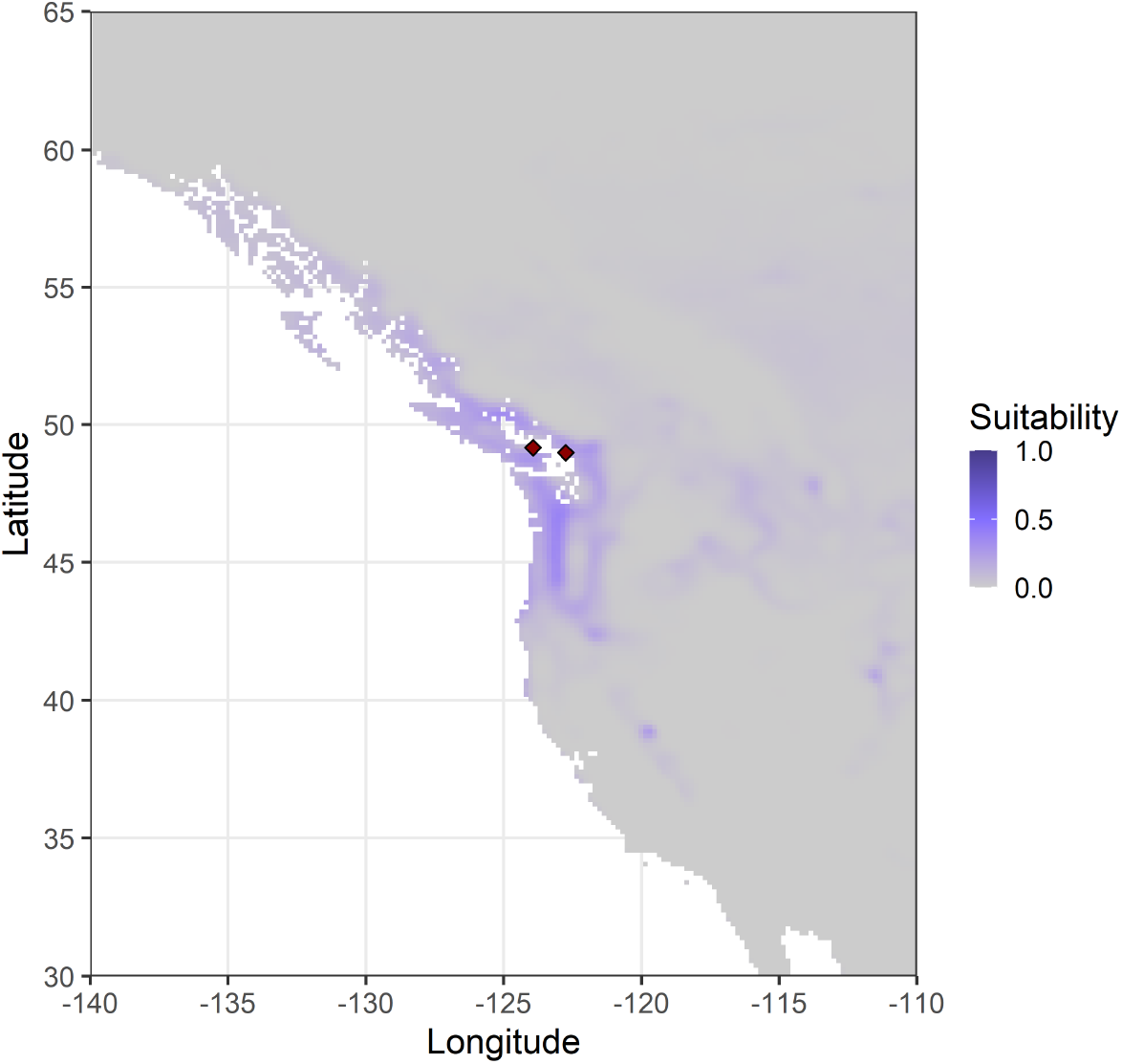
Suitability index obtained using the weighted-normal model for *V. mandarinia* plotted for the west coast of Canada and the United States. The red squares are confirmed occurrence points in each country.

